## supplementary Material for "AlphaFold2 and RoseTTAFold Predict Posttranslational modifications. Chromophore Formation in GFP-like Proteins" (from another version of this preprint: version 1, 2022-02-01)

**Supplementary Materials**

**Figure S1** Main chain i to i+4 "hydrogen bonding" alpha helical interactions measured and presented in Table S1.

**Figure S2** Alignment of all sequences used.

**Figure S3** Predicted possibility that structure forms a chromophore or cannot form a chromophore using LASSO model from Table S3 on AlphaFold2 data.

**Figure S4** RMSD overlap of the α-helix of the 1EMA-crystal with the α-helix of 1EMA as determined by RoseTTAFold for GFP-like proteins that will form a chromophore and those that do not.

**Figure S5** Predicted possibility that structure forms a chromophore or cannot form a chromophore using LASSO model from Table S5, which is based on RoseTTAFold data.

**Table S1** RMSD overlap of the α-helix of the 1EMA, 2AWJ, 1H4U crystal structures with the α-helix of the RoseTTAFold and AlphaFold determined structures, tight turn distances of RoseTTAFold and AlphaFold determined structures and the main chain i to i+4 "hydrogen bonding" alpha helical interactions. These are the geometric measurements that are crucial to chromophore formation. The structures are divided into two groups, GFP-like proteins that will form a chromophore and those that do not.

**Table S2** Description of all sequences used in this study.

**Table S3** Summary statistics of the RMSD overlap (in Angstrom) of the α-helix of the 1EMA-crystal with the α-helix of 1EMA as determined by AlphaFold2 for GFP-like proteins that will form a chromophore and those that do not.

**Table S4** LASSO model results for the alpha helix overlap with 1EMA-crystal as well as H-bond distances in angstrom between residues 61-65 (HD2), 62-66 (HD3), 70-74 (HD11) collected using AlphaFold2.

**Table S5** Summary statistics of the RMSD overlap (in Angstrom) of the α-helix of the 1EMA-crystal with the α-helix of 1EMA as determined by RoseTTAFold for GFP-like proteins that will form a chromophore and those that do not.

**Table S6** LASSO model results for the RMSD of the alpha helix overlap with 1EMA and 2AWJ-crystal structures, the tight turn distances, as well as H-bond distances in Angstrom between residues 61 and 65 (HD2), 62-66 (HD3), 63-67 (HD4), 64-68 (HD5) collected using RoseTTAFold simulations.

Figure S1: Main chain "hydrogen bonding" interactions measured and presented in Table S1. For an ideal $\alpha$-helix one expects to find hydrogen bonds between all i to i+1 residues (right). In the standard structures that have a chromophore the interactions shown in green were not formed because residues 65,66 and 67 were post-translationally modified.

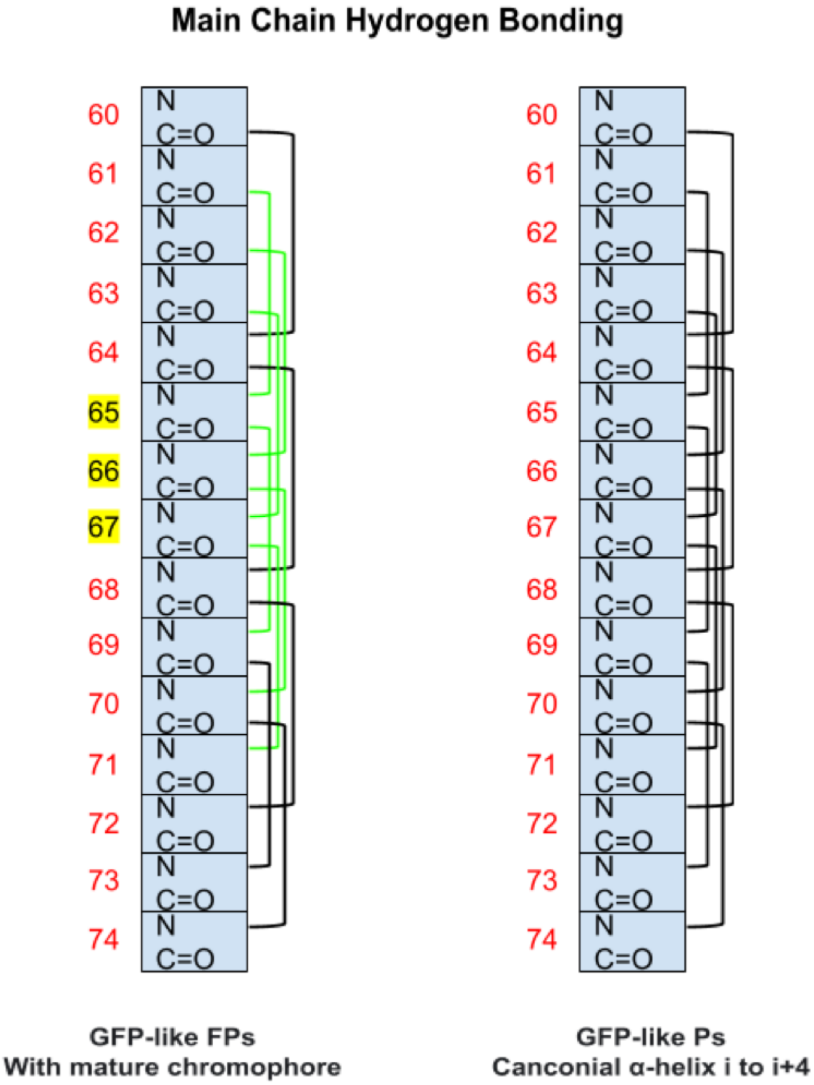

Figure S2: Alignment of all the sequences used in this paper. Shown relative to S65T GFP. Residues highlighted in yellow and blue make up the central helix. The 3 residues highlighted in blue correspond to the chromophore forming 65TYG67 in 1EMA. The origins of the GFP-like proteins are given in Table S2.

```
1EMA_1|Chain                DVN------G---HKFSVSGEGEGDATYGKLTLKFICT---T-GKLPVPWPTLVTTFTYG      67
1ZGO_1|Chains               TVN------G---HEFEIEGEGEGRPYEGHNTVKLKVT--KG-GPLPFAWDILSPQFQYG      68
1YHI_1|Chain                DVN------G---HKFSVSGEGEGDATYGKLTLKFICT---T-GKLPVPWPTLVTTFASG      68
2AWJ_1|Chain                DVN------G---HKFSVSGEGEGDATYGKLTLKFICT---T-GKLPVPWPTLVTTLTYG      68
1H4U_1|Chain                RIFVGSSQVPV--VFENTDLHSYVVMNHGRSYTAISTI---P-ETVGYSLLPLAPIGGII      98
6YLS_1|Chain                NVN------G---HHFVIDGDGTGKPYEGKQTMDLEVK--EG-GPLPFAFDILTTAFHYG      65
Dronpa-2                    AVN------G---HPFAIEGVGLGKPFEGKQSMDLKVK--EG-GPLPFAYDILTTVFCYG      64
PS-CFP                      DVN------G---HKFSVSGEGEGDATYGKLTLKFICT---T-GKLPVPWPTLVATLSYG      67
amFP495                     CVN------G---HYFTVKGEGNGKPYEGTQTSTFKVTMANG-GPLAFSFDILSTVFKYG      70
anm2CP                      VVN------D---QKFTIIADGSSKFPHGDFNVHAVCE---T-GKLPMSWKPICHLIQYG      67
P4                          DVN------G---HKFSVSGEGEGDATYGKLTLKFICT---T-GKLPVPWPTLVTTFSHG      67
AvicFP1                     DVH------G---HKFSVRGEGEGDAGSGKIEIKFVCT---T-GTLPVPWPTLVTTFCYG      67
cFP484                      NVN------G---HAFVIEGEGEGKPYDGTHTLNLEVK--EG-APLPFSYDILSNAFQYG      106
eYGFP                       NLN------G---EKFELVGGGVG--EEGRLEIEMKTK---D-KPLAFSPFLLSHCMGYG      57
dendFP                      NVN------G---HAFVIEGEGKGKPYEGTQTLNLTVK--EG-APLPFSYDILTTALHYG      64
dfGFP                       EIN------G---KKFKVAGEGFT-PNSGRFNMHAYCT---T-GDLPMSWVVIASPLQYG      66
DsRed-Express               SVN------G---HEFEIEGEGEGRPYEGTQTAKLKVT--KG-GPLPFAWDILSPQFQYG      68
EosFP                       NVN------G---HHFVIDGDGTGKPFEGKQSMDLEVK--EG-GPLPFAFDILTTAFHYG      64
eqFP578                     TVN------N---HHFKCTSEGERKPYEGTQTMKIKVV--EG-GPLPFAFDILATSFMYG      65
RFP611                      SVN------G---YQFKCTGEGDGNPYMGTQTMRIKVV--EG-GPLPFAFDVLATSFMYG      65
HcRed1-Blue                 TVN------G---HYFKCEGEGDGNPFAGTQSMRIHVT--EG-APLPFAFDILAPCCHYG      65
mKikGR                      SVN------G---HKFVITGKGSGRPYEGTQTVDLTVI--EG-GPLPFAFDILTTAFHYG      64
KO                          SVN------G---HEFTVEGEGTGKPYEGHQEMTLRVTMAKG-GPMPFSFDLVSHTFCYG      66
dLanYFP                     SFN------G---VDFDMVGRGTGNPNDGYEELNLKST---K-GDLQFSPWILVPQIGYG      70
Montipora                   TVN------G---HYFEVEGDGKGKPYEGEQTVKLTVT--KG-GPLPFAWDILSPLSQYG      64
mRed7                       SVN------G---HEFEIEGEGEGRPYEGTQTAKLKVT--KG-GPLPFAWDILSPQFMYG      69
ZYellow1                    CVN------G---HKFVITGEGIGYPFKGKQTINLCVI--EG-GPLPFSEDILSAGFKYG      68
B.fosteri                   DLG------D-DSDGYKITGEGLGQPLEGTQSLELNSA-----RDMPVNFNILGTIIQGN      72
B.forskali                  CFG------ED--HMVKVTGEGFSCPLEGAQTLELNST-----VPIPVNFNILGTLMQSN      71
B.infundibulum              HFE------DEPDSSYKVTGEGFAQPLDGCQNLELNCN-----SPLPINFNIVGTIIQTN      73
C.veneris                   HFE------DDPESPYKVTGEGFSQPLEGSQSLELNCA-----KPLPINFNILGTILQTN      73
D.kaloktenota               HFE------DDPESSYKVTGEGFSQPLEGSQSLELNCA-----RPLPINFNIVGTILQTN      73
D.glandiformis              EFQ------PG--TIIKVTGEGYSYAREGQQTLELNSI-----TSLPVNFGIVGTLIQTS      71
E.dunlapae                  VYE------D---QTYKLSGEGFGNPQEGQHTLEMRCF--GT-EACPLSWFVLGPVIQYS      72
H.rubra                     NFG------ED--ENVKITGEGFACPLEGSQTLELNSP-----KSLPINMNILGTIIQSN      71
Haeckelia_beehleri_FPL      QIT------EG--EDIKITGEGFSCPLEGSQNLELNSS-----TSLPINFNIVGTIIQSN      71
H.californensis             TFE------ED--LVIQVNGEGTASPREGVQTMELSCA--KY--AGPINLNIIGTIIQTN      73
P.bachei                    SYD------DE--LCVQVNGEGTGSPREGVQTMELSCP--NSPGPVAINLGIIGTIIQTN      75
Kiyohimea_sp                HFE------DDPESSYKVTGEGFSQPLEGSQSLELNCT-----RPLPINFNIVGTILQTN      73
L.lactea                    KID------D---MEYTITGEGFACPTEGQQNLELHCS--GT-SALPINWCILGTIIQCN      72
Lampea_sp                   KID------D---MEYTITGEGFACPTEGNQNLELHCS--GT-TALPINWCILGTIIQCN      72
L.cruentiventer             HFG------DE--HSYKVTGEGFSQPLDGAQTLALNCG-----KSMPVNFNIIGTIIQTN      71
L.pulchra                   HFE------DDPESSYKVTGEGFCQPMEGTQSLELNCT-----RSLPINLNIVGTIIQTN      73
Llyria_sp_B                 HFG------DE--HSYKVTGEGFSQPLDGAQSLELNCG-----KSMPINFNIIGTIIQTN      71
O.maculata                  HFG------DDPDMTFKVTGEGFAQPLEGSQSLELNCS-----TPLPINFNVVGTIIQTN      73
T.inconstans                SFG------DE--DTYKVTGEGFSQPLDGAQTLELNCG-----KSMPVNFNIVGTIIQTN      71
V.parallelum                HFE------DDEESPYKVTGEGFSQPLEGSQSLELNCT-----KPLPINFNVVGTLIQSV      73
XP_032228514.1:76-320       TVG------GL--SINSLFADTIGYSSDGRAYVATRGL---P-RPLAYSIRIVIPLASVV      92
XP_015751332.1:289-462      KIN------SV--DVPSINVDIIAMTKDGRAYVVIRDP---P-ASLQHSLKILTPFVNLV      92
KAB1270852.1:361-625        KVFVGSSPVPI--VFDNTDLHSYVVMNHGRSYTAISTI---P-ETVGYSLLPLAPIGGII      98
XP_012819171.1:387-646      KIYVGSNPVPV--VFEGIDLHSYVVVNDGRAYTAISTI---A-DSLGYSLLPLASIGGII      98
```

```
1EMA_1|Chain              VQCFSRYPDHMKRHDFFKS-AMPEGYVQERTIFFK-DDGNYKTRAEV--KFE---GDTLV    120
1ZGO_1|Chains             SKVYVKHPAD--IPDYKKL-SFPEGFKWERVMNFE-DGGVVTVTQDS--SLQ---DGCFI    119
1YHI_1|Chain              VQCFSRYPDHMKQHDFFKS-AMPEGYVQEATISFK-DDGNYKTRAEV--KFE---GDTLV    121
2AWJ_1|Chain              VQCFSRYPDHMKQHDFFKS-AMPEGYVQEMTISFK-DDGNYKTRAEV--KFE---GDTLV    121
1H4U_1|Chain              GWMFAVEQD--GFKNGF-S-ITGGEFTRQAEVTFLGHPGKLVLKQQFSGIDEHG-HLTIS    153
6YLS_1|Chain              NRVFVKYPDN--IQDYFKQ-SFPKGYSWERSLTFE-DGGICNARNDI--TME---GDTFY    116
Dronpa-2                  NRVFAKYPEN--IVDYFKQ-SFPEGYSWERSMNYE-DGGICNATNDI--TLD---GDCYI    115
PS-CFP                    VQCFSRYPDHMKQHDFFKS-AMPEGYIQERTIFFE-DDGNYKSRAEV--KFE---GDTLV    120
amFP495                   NRCFTAYPTS--MPDYFKQ-AFPDGMSYERTFTYE-DGGVATASWEI--SLK---GNCFE    121
anm2CP                    EPFFAKYPNG--ISHFAQE-CFPEGLTIDRTVRFE-NDGTMTSHHTY--ELD---GTCVI    118
P4                        VQCFSRYPDHMKQHDFFKS-AMPEGYVQERTIFFK-DDGNYKTRAEV--KFE---GDTLV    120
AvicFP1                   VQCFTRYPEHMKQHDFYKS-AMPDGYIQKRTISFQ-DDGHYKTRAEV--KFE---GDTLV    120
cFP484                    NRALTKYPDD--IADYFKQ-SFPEGYSWERTMTFE-DKGIVKVKSDI--SME---EDSFI    157
eYGFP                     FYHFASFPKG--TKNIYLHAATNGGYTNTRKEIYE-DGGILEVNFRY--TYE---FNKII    109
dendFP                    NRVFTKYPED--IPDYFKQ-SFPEGYSWERTMTYE-DKGICTIRSDI--SLE---GDCFF    115
dfGFP                     FHMFAHYPED--ITHFFQE-CFPGSYTLDRTLRME-GDGTLTTHHEY--SLK---DGCVT    117
DsRed-Express             SKVYVKHPAD--IPDYKKL-SFPEGFKWERVMNFE-DGGVVTVTQDS--SLQ---DGSFI    119
EosFP                     NRVFAEYPDH--IQDYFKQ-SFPKGYSWERSLTFE-DGGICIARNDI--TME---GDTFY    115
eqFP578                   SKTFINHTQG--IPDLFKQ-SFPEGFTWERITTYE-DGGVLTATQDT--SLQ---NGCII    116
RFP611                    SKTFIKHTKG--IPDFFKQ-SFPEGFTWERVTRYE-DGGVITVMQDT--SLE---DGCLV    116
HcRed1-Blue               SKTFVHHTAE--IPDFWKQ-SFPEGFTWERTTTYE-DGGILTAHQDT--SLE---GNCLI    116
mKikGR                    NRVFVEYPEE--IVDYFKQ-SFPEGYSWERSMSYE-DGGICLATNNI--TMKKDGSNTFV    118
KO                        HRPFTKYPEE--IPDYFKQ-AFPEGLSWERSLQFE-DGGFAAVSAHI--SLR---GNCFE    117
dLanYFP                   FHQYLPFPDG--MSPFQAAMKDGSGYQVHRTMQFE-DGASLTSNYRY--TYE---GSHIK    122
Montipora                 SIPFTKYPED--IPDYVKQ-SFPEGYTWERIMHFE-DGAVCTVSNDS--SIQ---GNCFI    115
mRed7                     SKAYVKHPAD--IPDYFKL-SFPEGFKWERVMNFE-DGGAVTVTQDS--SLE---DGTLI    120
ZYellow1                  DRIFTEYPQD--IVDYFKN-SCPAGYTWGRSFLFE-DGAVCICNVDITVSVK---ENCIY    121
B.fosteri                 LRMFTQYTGT-MVYDFFKT-CFPGTMTTEFTGTFS-DGVKLEGCCTI--THV---KDTLI    124
B.forskali                FRMFVQYLGI-TAYDFFKT-AFPGTMVVEMEGSFS-DGLTVRGTFTL--SYV---KDTLI    123
B.infundibulum            FRMFTQYTGT-MVYDFFKT-GFPGSMNVELEGKFT-DGMTVRATCTL--THV---KETLI    125
C.veneris                 FRMNTQYSGT-MVYDFFKT-SFPGTMTVEMDGKFC-DDMTVRATCTL--TYM---KDHLI    125
D.kaloktenota             FRMFTQYTGT-MVYDFFKT-SFPGTMTVEMEGKFC-DDMTVRATCTL--THV---KDTLI    125
D.glandiformis            FRMYTLYPGN-MIYDFFKT-SFPGTMNVEMKGVFS-DGMTMDSVCTL--SFV---NGTLI    123
E.dunlapae                YRMFTQYSGN-GMYDFFKT-SFPGGLSTESVCTFN-DGATITGSHNI--SFV---KDIVV    124
H.rubra                   FRMFIQYRGS-SVYDFFKT-SFPGTMVVETEGDFS-DGTSVRGTSTL--TYV---KEALI    123
Haeckelia_beehleri_FPL    FRMFTQYTGN-TVYDFFKT-AFPGTMVVELEGSFC-DGLSLKGSCTL--TYV---KDALI    123
H.californensis           LRAFTGYTGS-NLYDFFKT-SFPGSPKTEIEAMFV-DGAIVSGSCTL--SFV---KSTLM    125
P.bachei                  LKPFVNYMGS-QVYDFYKT-NFPGSPTTDIEAIFV-DGMTISGSSTL--SFV---KNILM    127
Kiyohimea_sp              FRMFTQYTGT-MVYDFFKT-SFPGTMTVEMEGKFC-DDMTVRVTCTL--THV---KDTLI    125
L.lactea                  FKLFTQYKGS-NEYDFFKT-SFPGGLKVEYVGSFV-DGAEVSGNSTM--SYV---KDTVI    124
Lampea_sp                 FKLFTQYKGS-NEYDFFKT-SFPGGLKVEYVGSFV-DGAEVSGNSTM--SYV---KDTVI    124
L.cruentiventer           FRMFTQYTGT-MMYDFFKT-SFPGTMTIDMEGRFS-DGLVLTATCTL--TYV---KDTLI    123
L.pulchra                 FRMFTQYTGS-MVYDFFKT-SFPGTMTIESDGKFC-DDMTVRTSCTI--TLV---KDAFI    125
Llyria_sp_B               FRMFTKYTGA-MVYDFFKT-SFPGTMTIEMDGSFS-DGLVLKATCTL--TYV---KDTLI    123
O.maculata                FRMFTQYTGS-MVYDFFKT-SFPGQMNVEMDGTFT-DELTVKVSCTL--TYV---KDCLI    125
T.inconstans              FRMFTQYTGT-TVYDFFKT-SFPGTMTVDMEGSFS-DGLGFKANCTL--TYV---KDTLI    123
V.parallelum              FRMNTQYSGT-MVYDFFKT-SFPGTMNVEMEGTFC-DEMTVKATCTL--TYM---KDHLI    125
XP_032228514.1:76-320     GWMFGMPVHK-GLKNGY-A-VVGDRFTRNVTVVFD-SGEMLSIDQQFQGKDSSSPHVRVQ    148
XP_015751332.1:289-462    GWMFAVKKRM-AVRNGI-Q-VIGEKFTRSAIISFN-SGEALYLNQRFTGKDATGSYISIV    148
KAB1270852.1:361-625      GWMFAVEQG--GFKNGF-S-ITGGEFTRLAEVTFVGHPDKLVIKQRFSGIDEHG-HLTID    153
XP_012819171.1:387-646    GWMFSVQQP--GYKNGF-S-ITGGEFTRRAEVTFLQSNKKLVILQKFSGIDEHG-HLTIS    153
```

Figure S3: Prediction results using LASSO model from Table S4 on AlphaFold2 data. GFP-like proteins that will form a chromophore and those that do not.

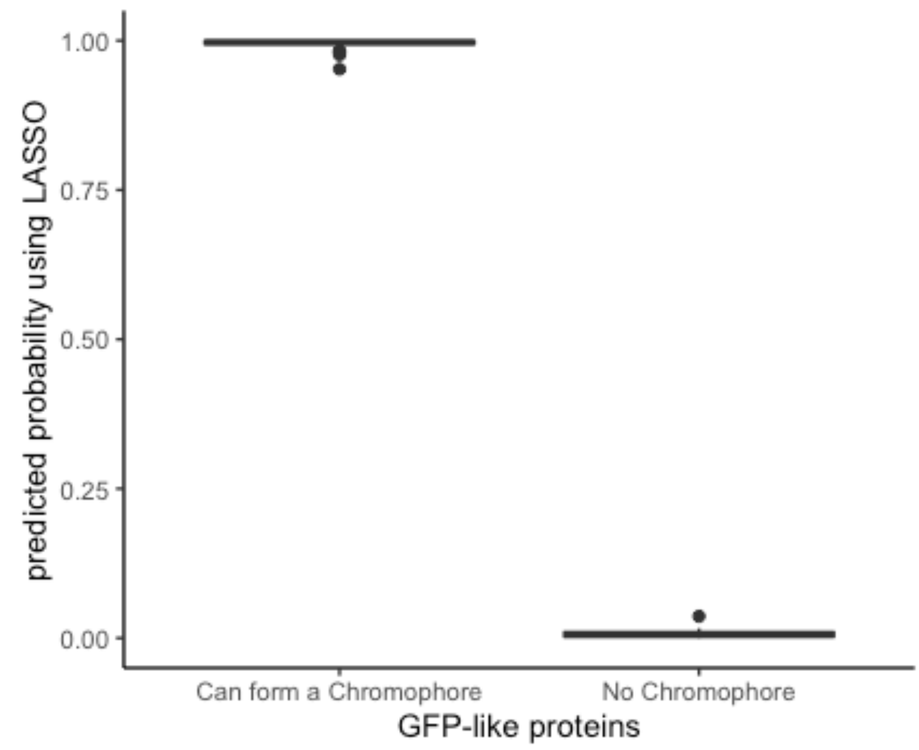

Figure S4: RMSD overlap of the α-helix of the 1EMA-crystal with the α-helix of 1EMA as determined by RoseTTAFold for GFP-like proteins that will form a chromophore and those that do not.

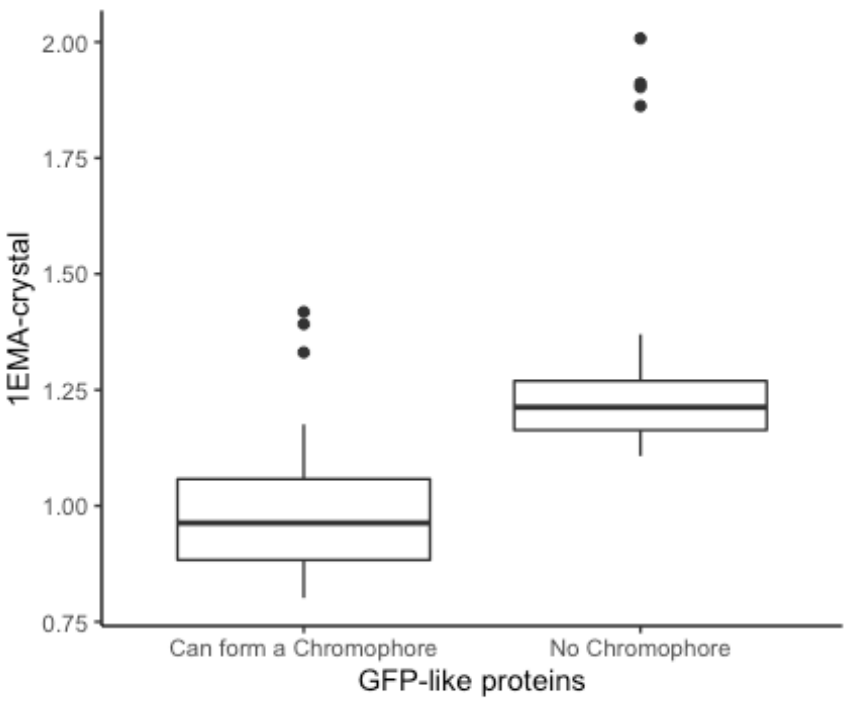

Figure S5: Predicted possibility that structure forms a chromophore or cannot form a chromophore using LASSO model from Table S6, which is based on RoseTTAFold data.

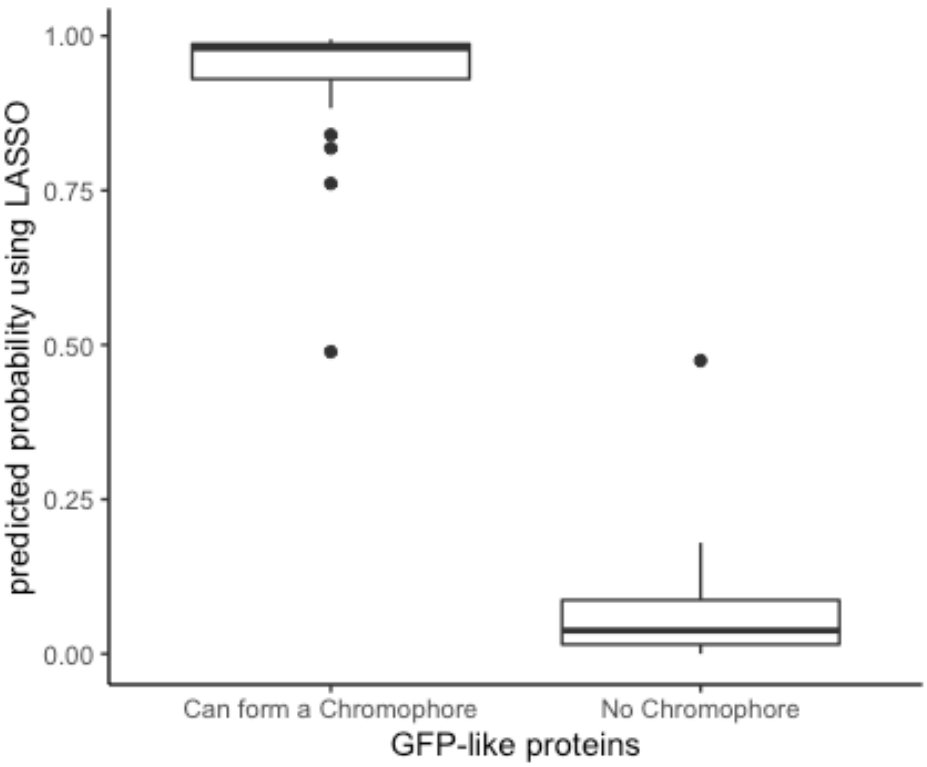

Table S2: Description of all sequences used in this study.

| | Name | Percent Identity to 1EMA | Species | Source |
|---|---|---|---|---|
| **Standards** | 1EMA[15] | 100 | *Aequorea victoria* | rcsb.org |
| | 1YHI[27] | 97 | *Aequorea victoria* | |
| | 2AWJ[16] | 97 | *Aequorea victoria* | |
| | 1ZGO[28] | 23 | *Discosoma* | |
| | 6YLS[29] | 25 | *Lobophyllia hemprichii* | |
| | 1H4U[17] | 17 | *Mus musculus* | |
| **Capable of forming chromophore** | Dronpa-2[30] | 24 | *Echinophyllia sp. SC22* | https://www.fpbase.org/lineage/ |
| | PS-CFP[31] | 89 | *Aequorea coerulescens* | |
| | amFP495[32] | 25 | *Anemonia majano* | |
| | anm2CP[33] | 25 | *Anthoathecata* | |
| | P4[34] | 98 | *Aequorea victoria* | |
| | AvicFP1[35] | 80 | *Aequorea victoria* | |
| | cFP484[36] | 26 | *Clavularia sp.* | |
| | eYGFP[37] | 17 | *Chiridius poppei* | |
| | dendFP[38] | 26 | *Dendronephthya sp.* | |
| | dfGFP[39] | 28 | *Olindias formosus* | |
| | DsRed-Express[40] | 23 | *Discosoma sp.* | |
| | EosFP[41] | 26 | *Lobophyllia hemprichii* | |
| | eqFP578[42] | 19 | *Entacmaea quadricolor* | |
| | RFP611[43] | 21 | *Entacmaea quadricolor* | |
| | HcRed1-Blue[44] | 20 | *Heteractis crispa* | |
| | mKikGR[45] | 27 | *Favia favus* | |
| | KO[46] | 23 | *Verrillofungia concinna* | |
| | dLanYFP[47] | 24 | *Branchiostoma lanceolatum* | |
| | *Montipora sp.* #20[48] | 23 | *Montipora sp.* #20 | |
| | mRed7[49] | 30 | synthetic | |
| | zYellow1 | 22 | *Zoanthus sp.* | |
| **Incapable of forming chromophore** | *B. forskali[19]* | 20 | *B. forskali[19]* | Non-excitable fluorescent protein orthologs found in ctenophores [19] |
| | *B. fosteri[19]* | 20 | *B. fosteri[19]* | |
| | *B. infundibulum[19]* | 18 | *B. infundibulum[19]* | |
| | *C. veneris[19]* | 20 | *C. veneris[19]* | |
| | *D. glandiformis[19]* | 23 | *D. glandiformis[19]* | |
| | *D. kaloktenota[19]* | 20 | *D. kaloktenota[19]* | |
| | *E. dunlapae[19]* | 22 | *E. dunlapae[19]* | |
| | *H. californensis[19]* | 18 | *H. californensis[19]* | |
| | *H. rubra[19]* | 19 | *H. rubra[19]* | |
| | *H. beehleri[19]* | 19 | *H. beehleri[19]* | |
| | *Kiyohimea[19]* | 19 | *Kiyohimea[19]* | |

| | | | | |
|---|---|---|---|---|
| | *L. cruentiventer[19]* | 21 | *L. cruentiventer[19]* | |
| | *L. lactea[19]* | 20 | *L. lactea[19]* | |
| | *L. pulchra[19]* | 18 | *L. pulchra[19]* | |
| | *Lampea[19]* | 20 | *Lampea[19]* | |
| | *Llyria[19]* | 21 | *Llyria[19]* | |
| | *O. maculata[19]* | 19 | *O. maculata[19]* | |
| | *P. bachei[19]* | 17 | *P. bachei[19]* | |
| | *T. inconstans[19]* | 20 | *T. inconstans[19]* | |
| | *V. parallelum[19]* | 18 | *V. parallelum[19]* | |
| | KAB1270852 | 18 | *Camelus dromedarius* | https://blast.ncbi.nlm.nih.gov/Blast.cgi?PAGE=Proteins |
| | XP012819171 | 17 | *Xenopus tropicalis* | |
| | XP032228514 | 12 | *Nematostella vectensis* | |

Table S3: Summary statistics of the RMSD overlap (in Angstrom) of the $\alpha$-helix of the 1EMA-crystal with the $\alpha$-helix of 1EMA as determined by AlphaFold2 for GFP-like proteins that will form a chromophore and those that do not.

| Groups | Minimum | First Quartile | Median | Mean | Third Quartile | Maximum |
|---|---|---|---|---|---|---|
| Does not form chromophore | 1.165 | 1.244 | 1.274 | 1.372 | 1.320 | 2.086 |
| Forms chromophore | 0.574 | 0.734 | 0.784 | 0.800 | 0.860 | 1.049 |

Table S4: LASSO model results for the alpha helix overlap with 1EMA-crystal as well as H-bond distances in Angstrom between residues 61-65 (HD2), 62-66 (HD3), 70-74 (HD11) collected using AlphaFold2.

| Variable names | coefficients | exp(coefficients) |
|---|---|---|
| 1EMA-crystal | -7.284 | 0.001 |
| HD2 | 0.395 | 1.485 |
| HD3 | 0.745 | 2.107 |
| HD11 | 0.788 | 2.199 |

From the model results (as listed above in Table S4) The first column listed the variables that are selected out, the second column are the coefficients estimates, and the third column are the exponentiated coefficients. Each exponentiated coefficient is the change in odds in the multiplicative scale for a unit increase in the corresponding predictor variable holding other variables at certain value. For example, the exponentiated coefficient for 1EMA-crystal is 0.001, meaning that holding the other variables at a fixed value, with one unit increase in the alpha helix overlap of 1EMA-crystal, the odds of being GFP-like proteins that will form a chromophore will decrease by 99.9% (=100%-0.1%). Also, from the boxplot of the predictions of the LASSO model against the two groups, we can see there is a clear distinction.

Table S5: Summary statistics of the RMSD overlap (in Angstrom) of the $\alpha$-helix of the 1EMA-crystal with the $\alpha$-helix of 1EMA as determined by RoseTTAFold for GFP-like proteins that will form a chromophore and those that do not.

| Groups | Minimum | First Quartile | Median | Mean | Third Quartile | Maximum |
|---|---|---|---|---|---|---|
| Do Not Form Chromophore | 1.1074 | 1.16335 | 1.21275 | 1.317071 | 1.269375 | 2.0078 |
| Will Form Chromophore | 0.8014 | 0.8834 | 0.963 | 1.003562 | 1.057825 | 1.4181 |

Table S6: LASSO model results for the RMSD of the alpha helix overlap with 1EMA and 2AWJ-crystal structures, the tight turn distances, as well as H-bond distances in Angstrom between residues 61 and 65 (HD2), 62-66 (HD3), 63-67 (HD4), 64-68 (HD5) collected using RoseTTAFold simulations.

| Variable names | coefficients | exp(coefficients) |
|---|---|---|
| 1EMA-crystal | -3.054 | 0.047 |
| 2AWJ-crystal | -0.534 | 0.586 |
| tight_turn_D | 0.721 | 2.057 |
| HD2 | 1.064 | 2.897 |
| HD3 | 1.241 | 3.459 |
| HD4 | 0.601 | 1.824 |
| HD5 | 0.541 | 1.718 |

The first column listed the variables that are selected out, the second column are the coefficients estimates, and the third column are the exponentiated coefficients. Each exponentiated coefficient is the change in odds in the multiplicative scale for a unit increase in the corresponding predictor variable holding other variables at certain value. For example, the exponentiated coefficient for 1EMA-crystal is 0.078, meaning that holding the other variables at a fixed value, with one unit increase in the alpha helix overlap of 1EMA-crystal, the odds of being GFP-like proteins that will form a chromophore will decrease by 95.3% (=100%-4.7%).
